# Transient network synchrony of the developing medial entorhinal cortex

**DOI:** 10.1101/121459

**Authors:** Julia Dawitz, J.J. Johannes Hjorth, Tim Kroon, Marta Ruiperez-Alonso, Nandakumar Chandrasekhar, Henrike Hartung, Ileana L. Hanganu-Opatz, Huibert D. Mansvelder, Rhiannon M. Meredith

## Abstract

The medial entorhinal cortex (MEC) contains a variety of specialized spatially-tuned neurons whose properties emerge during the third postnatal week onwards in rodents. How neuronal networks underlying the spatial firing patterns are formed is largely unknown but they are hypothesized to develop from topographic modules of synchronized neurons in superficial MEC. Here, we show that developing MEC neuronal networks in the second postnatal week are synchronously active in spatially-grouped modules. Network synchrony is intrinsic to MEC and desynchronized just prior to the emergence of spatially-tuned firing properties. The MEC network is modulated but not driven by the immature hippocampus and is tightly-coupled to neighboring neocortical networks. Unlike hippocampal networks, developing modules are dominated by glutamatergic excitation rather than GABAergic inhibition. Our results demonstrate that intrinsically synchronous modules exist in immature MEC: these may play a key role in establishing and organizing circuitry necessary for spatially-tuned firing properties of MEC neurons.

---

The medial entorhinal cortex (MEC) contains specialized cell types whose firing is tuned to aspects of an animal’s position and orientation in the environment, constituting a neuronal representation of space (Moser and Moser 2008). The spatially-tuned firing properties of head-direction, grid, boundary and conjunctive cells emerge during the third postnatal week of development in rodents (Langston, Ainge et al. 2010, Wills and Cacucci 2014). In young adult rodents, superficial MEC layers are modulated by recurrent inhibition (Couey, Witoelar et al. 2013, Pastoll, Solanka et al. 2013) and grid cell firing patterns arise from local inhibitory connectivity in attractor network models (McNaughton, Battaglia et al. 2006, Couey, Witoelar et al. 2013, Pastoll, Solanka et al. 2013). However, it is not known how the network forms prior to spatial exploration and eye-opening in rodent pups. From computational modelling, it is hypothesized that topographic activity modules exist in immature MEC and that synchronous waves of coordinated activity could underlie the formation of such a spatial network (McNaughton, Battaglia et al. 2006).

In adulthood, the MEC is reciprocally connected to neocortical and parahippocampal regions, including the hippocampal formation (van Strien, Cappaert et al. 2009). Grid cell activity in MEC is shaped by excitatory hippocampal projections in the mature brain (Bonnevie, Dunn et al. 2013) and CA1 place cell fields are less precise following MEC layer III ablation (Brun, Leutgeb et al. 2008). Thus, in mature rodents recurrent connections exist between activity in MEC and neighboring hippocampal and neocortical regions. Immature hippocampus and neocortex are characterized by synchronized patterns of network activity. Synchronous network activity is restricted to specific developmental periods (Blankenship and Feller 2010) and essential for structural and functional maturation including axonal growth, synapse plasticity and sensory map formation (Blankenship and Feller 2010) (Spitzer 2006). Slow synchronous bursts of activity, so-called giant depolarizing potentials, are mediated by GABA in the immature hippocampus and thought to be required for the development of hippocampal network connectivity (Ben-Ari 2001). Therefore, we tested experimentally whether the theorised network modules exist in immature MEC and if so, at what developmental stages. Further, given the proximity and strong connectivity of MEC with the hippocampus at maturity, we determined whether network synchrony is driven by hippocampal inputs and is mediated via GABAergic mechanisms.

Using in vivo extracellular recordings, in vitro multiphoton calcium imaging, multi-electrode and paired cell recordings, we tested the three hypotheses that (i) topographic modules exist during early development, (ii) the hippocampus drives MEC network synchrony and that (iii) this is mediated via GABAergic transmission. We determined that synchronous modules of activity are intrinsic to the early development of MEC networks in the rodent during a restricted developmental period prior to eye-opening and spatial navigation. In contrast to mature MEC networks and immature hippocampal networks (Bonnevie, Dunn et al. 2013) (Couey, Witoelar et al. 2013) (Pastoll, Solanka et al. 2013), our data reveal highly synchronous glutamatergic activation *in vivo* and *in vitro* that is intrinsically-generated across layers. Immature MEC networks synchronize and modulate neighboring neocortex validating previous speculation(Garaschuk, Linn et al. 2000) but are neither driven by hippocampal nor neocortical activity. These modular networks, predicted by a computational model for path integration (McNaughton, Battaglia et al. 2006), may establish and tune the MEC circuitry to enable emergence of spatially-tuned cell properties that play a key role in spatial processing and navigation (Wills and Cacucci 2014).

## Results

### Spontaneous bursts of activity in developing medial entorhinal cortex

Discontinuous oscillatory activity is a hallmark pattern during early network development of many brain regions (Blankenship and Feller 2010). To determine whether MEC exhibits such rhythmic network activity, we performed extracellular recordings of the local field potential (LFP) and multiple unit activity (MUA) in neonatal [postnatal day (P) 7-8] urethane-anesthetized rodent pups (Fig.1a, n=4 pups). Neonatal MEC showed discontinuous patterns of oscillatory activity with a relatively high occurrence (P7-8: 0.12±0.01Hz, mean±SEM, Fig.1b,d) when compared to neocortical regions (Yang, Hanganu-Opatz et al. 2009). Oscillatory bursts were of short duration and large amplitude (3.3±0.2s, 249±7.3μV, n=160 events) and contained a predominant theta band that was phase-locked to high-frequency MUA (Fig. 1c, 6.2±0.1Hz, n=160 events). Additionally, higher frequency components in the gamma band range were detected.

**Figure 1:**
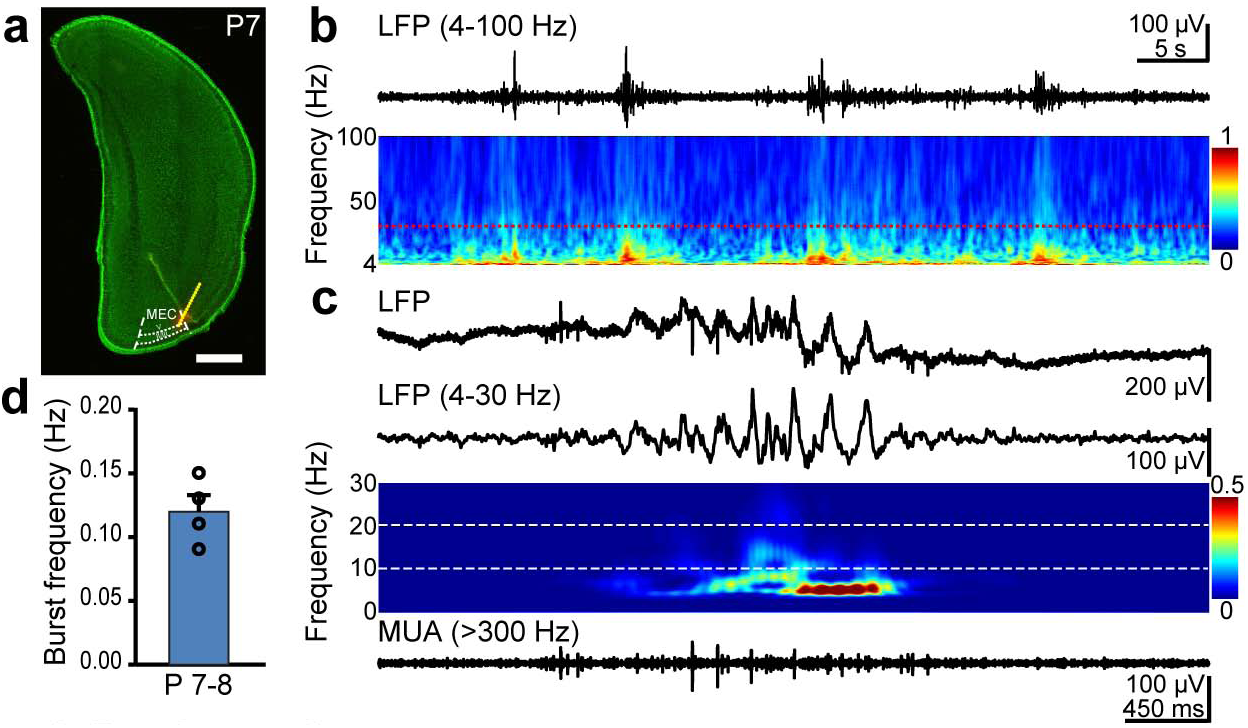
Patterns of network activity in the neonatal MEC *in vivo*. (**a**) Digital photomontage reconstructing the location of the DiI-labeled recording electrode (orange) in the MEC of a Nissl-stained 100μm-thick coronal section (green) from a P7 pup. Yellow dots mark the recording sites covering the entorhinal layer II/III. Scale bar 1mm. (**b**) Band-pass filtered (4-100Hz) extracellular field potential recording of discontinuous oscillatory activity in the MEC accompanied by the color-coded wavelet spectra of the LFP at identical time scale. The red dotted line marks the lower border of gamma frequency band (30Hz). (**c**) Characteristic theta burst displayed before (*top*) and after band-pass (4-30Hz) filtering (*middle*) and the corresponding MUA after 200Hz high-pass filtering (*bottom*). Color-coded frequency plot shows the wavelet spectrum of LFP at identical time scale. (**d**) Occurrence of MEC theta bursts (mean ± SEM) with superimposed values for individual animals (n=4).

To unravel the mechanisms underlying the network bursts we utilized an *in vitro* slice approach. Field recordings in horizontal slices of superficial layers of MEC (sMEC) confirmed the occurrence of network bursts with activity patterns similar to *in vivo* (unpaired t-test, t(25)=-1.5, p=0.15, Fig. 1d, Suppl.Fig.1c). Using multiphoton calcium imaging for single cell resolution we found that networks were comprised of silent and active cells, of which the majority was synchronously active (Fig.2a). To establish the developmental time-course of activity we mapped spontaneous network events during the second postnatal week, prior to the onset of spatially-tuned neuronal firing properties in MEC (Langston, Ainge et al. 2010, Wills and Cacucci 2014). In each neuron somatic calcium events reflecting suprathreshold activity were binarized and their synchronization across the network was calculated (Fig.2b-d, see Methods). Network activity peaked at P10/11 with the highest proportion of active neurons (40.1±4.9% cells) and significantly decreased towards P14/15 (7±2% cells, Fig.2c, blue bars). Within the active network, the fraction of active neurons that were synchronized followed a similar profile, peaking at P10/11 (31±4% cells) and decreasing by P14/15 (2±1% cells, Fig.2c, black bars). The frequency of active neurons was strongly developmentally regulated being higher at P10/11 (0.055±0.007Hz) than at P14/15 (0.008±0.004Hz). Frequency of synchronized bursts of events across the network (‘network events’ -see Methods for definition), also peaked at P10/11 (0.050±0.013Hz) and virtually disappeared by P14/15 (0.002±0.001Hz, Fig.2d). Despite a decrease in activity of all neurons with age, the decrease was more prominent for synchronized network events, changing significantly from comprising 79.3±11.1% of all activity at P10/11 to 29.6±15.5% at P14/15. Unless otherwise stated, all data in the following paragraphs are reported from peak activity ages P8-11.

**Figure 2:**
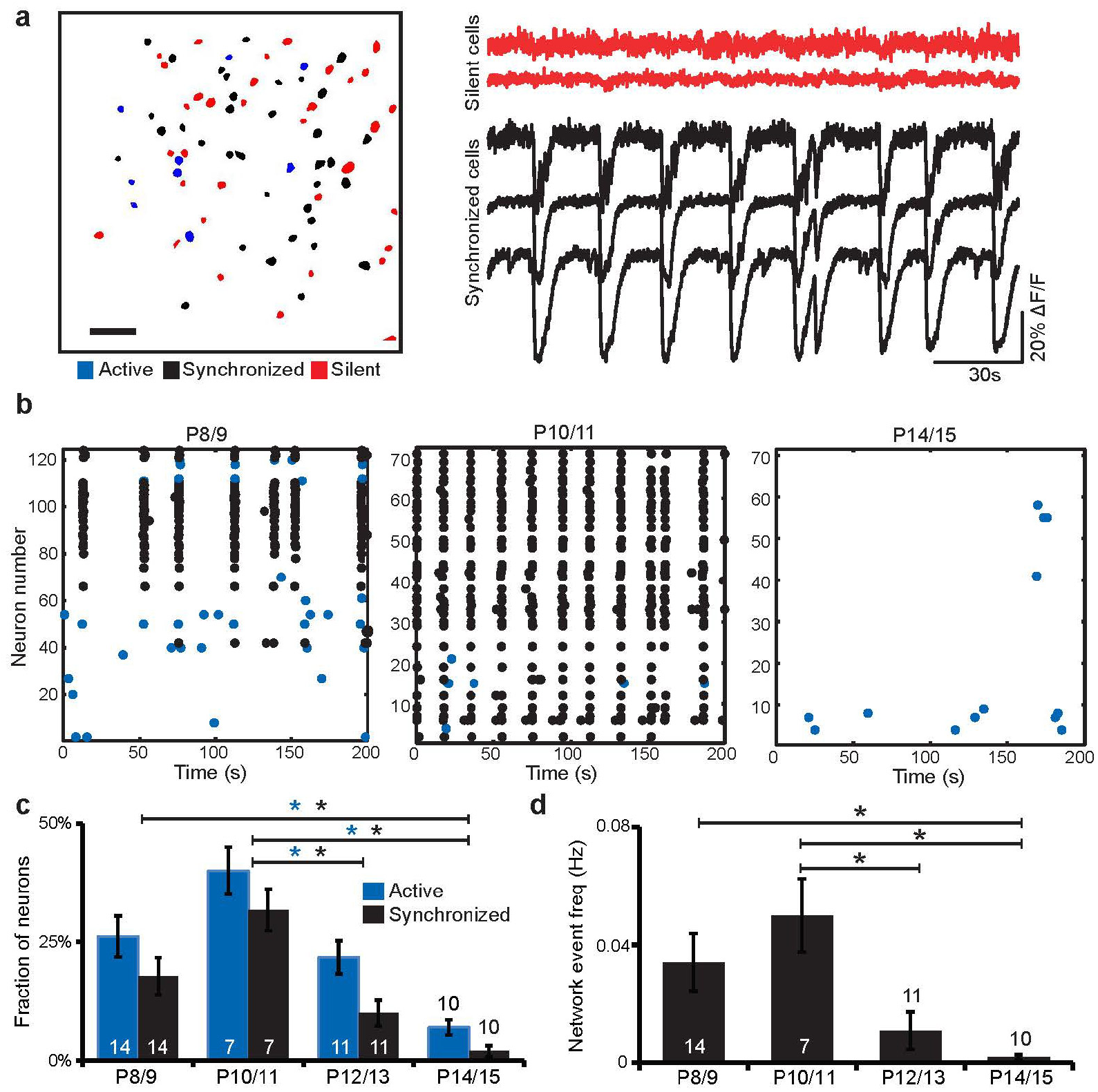
Developmental dynamics of spontaneous sMEC activity in vitro. (**a**) Contour map of Fura2-AM ester bulk-loaded cells in superficial medial entorhinal cortex networks imaged at P10/11. Active neurons (blue), silent neurons (red) and synchronized neurons (black) indicated. *Scalebar 50μm*. Example traces of silent and synchronously active neurons. (**b**) Representative raster plots of sMEC network activity at P8/9, P10/11, and P14/15, color coded as in a. (**c**) Quantification of the fraction of active and synchronized neurons during postnatal development (*n*(*networks imaged*)=*14, 7, 11, 10 respectively at P8/9, P10/11, P12/13 and P14/15, indicated in individual bars*) showed a significant age-dependent peak in activity (Bonferroni corrected, P8/9 vs. P14/15, p<0.001, and P10/11 vs. both P12/13, p<0.03, and vs. P14/15, p<0.001, one-way ANOVA, F(3,41)=10.1) and synchronization (Bonferroni corrected, P8/9 vs. P14/15 p<0.01, P10/11 vs. P12/13 p=0.001 and P10/11 vs. P14/15, p<0.001, one-way ANOVA, F(3,41)=11.1) at P10/11. (**d**) Frequency of network events (see *Methods* for definition) is highest at P10/11 and virtually absent by P14/15 (Bonferroni corrected: P8/9 vs. P14/15, p<0.05, P10/11 vs. P12/13, p<0.05, and P10/11 vs. P14/15, p<0.01, one-way ANOVA, F(3,41)=5.8).

### Configuration of synchronous network modules

In adult rodents, grid cells are the predominant spatially-tuned cell type found in layer II sMEC (Sargolini, Fyhn et al. 2006) and of these, the majority are stellate cells (Domnisoru, Kinkhabwala et al. 2013, Schmidt-Hieber and Hausser 2013). Simultaneous cell-attached recordings of stellate cells with calcium imaging (Fig.3a *left*) confirmed that immature stellate cells fire in synchrony with the network and thus, constitute a main component of the developmental bursts observed in sMEC (Fig.3a, Suppl Fig1). To test whether synchronized neurons were not only functionally- but also spatially-clustered in modules, as proposed by a theoretical model for grid cell development (McNaughton, Battaglia et al. 2006), we investigated their physical distribution across sMEC (Fig.3b, Suppl.Fig.2). Synchronized neurons were homogenously distributed with similar density across both superficial layers (II and III) (Fig.3b). However, the average distance between a synchronized neuron and its synchronized neighbors was significantly shorter compared to its non-synchronized neighbors (median: 156μm synchronized vs. 196μm non-synchronized, Fig.3c *left, middle*). Further investigation using a Sholl-like analysis revealed a distance-dependent distribution profile with the greatest number of synchronized neighbors grouped 50-100μm away (mean 4.54 neighbouring cells) and those of non-synchronized neighbors at 150-200μm (mean 10.93 neighbouring cells) (Fig.3c *right*). A comparison of these spatial patterns revealed a significantly different, non-random distribution of synchronized compared to nonsynchronized neighboring neurons within the stained networks (red line, Fig.3c *right*).

**Figure 3:**
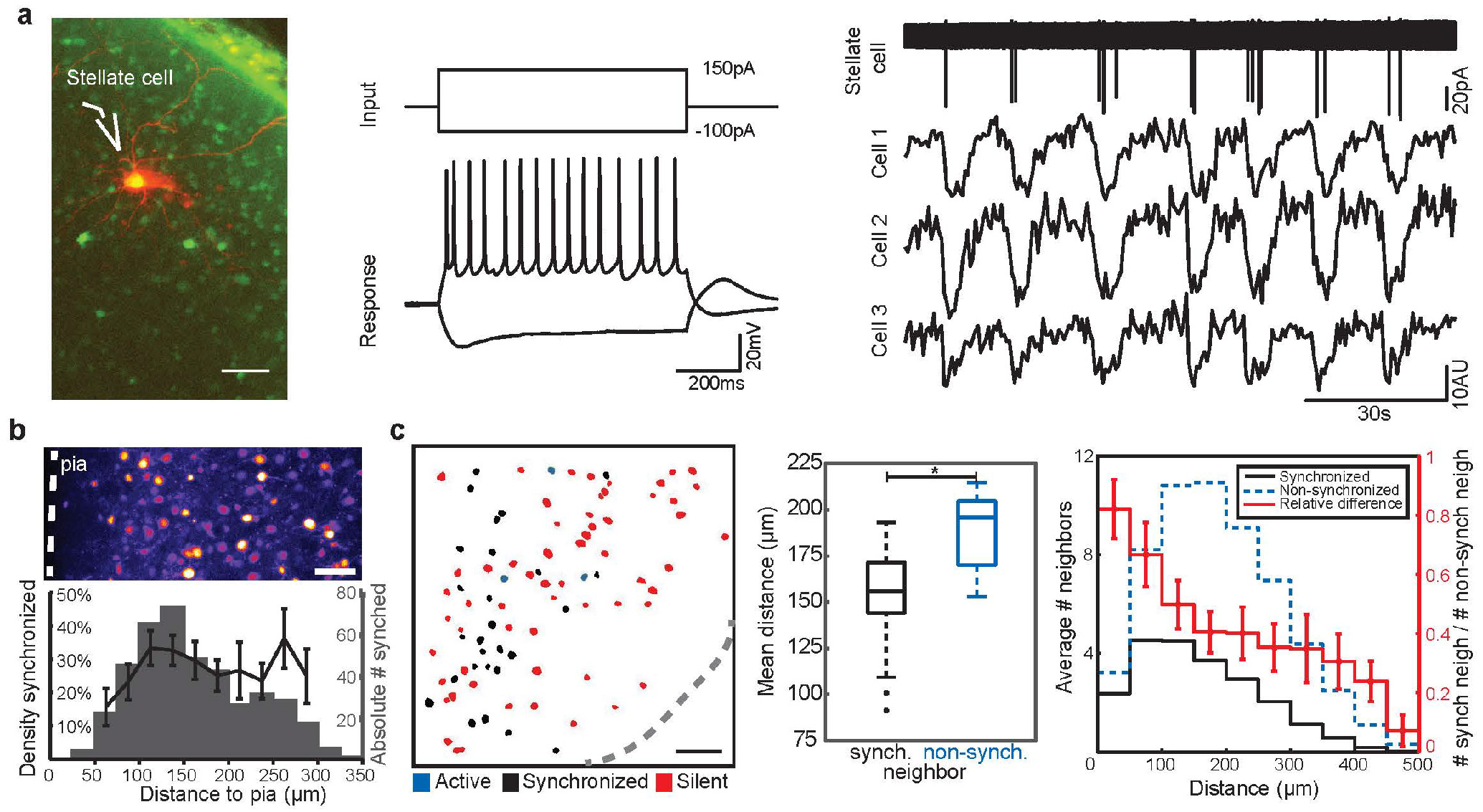
Cellular correlates of sMEC activity and spatial distribution of synchronized neurons. (**a**) Fluorescent image (*left*) of patched neuron (red) and Fura2-AM ester bulk-loaded neurons (green) with corresponding spike profile (*middle*). Parallel cell-attached and calcium recordings (*right*) show that identified stellate cells spike in synchrony with the network. *Scale bar 50μm*. (**b**) Two-photon image of sMEC network loaded with Fura2-AM ester dye. Dashed line indicates pia. Scale bar 50μm (top). *Bottom*: Proportion of synchronized neurons relative to total labeled neurons (density, black line) does not differ depending on distances across superficial layers of MEC relative to the pia (one-way ANOVA, F(9,153)=0.99, p=0.45). Total number of detected synchronized neurons per distance bin (gray bars). *N*=*18 slices*. (**c**) Example contour map illustrating non-random spatial distribution of synchronized neurons (black). *Scale bar 50μm* (*left*). Mean distances between synchronized neurons are significantly lower than between non-synchronized neurons across the entire network (Wilcoxon Signed Rank test, two-sided, W=3.72, p<0.0001). *N*=*18 slices* (*middle*). Spatial distribution of synchronized neurons and nonsynchronized (active or silent) neurons throughout the superficial network, indicating mean number of neighboring neurons (*lefthand axis*) per 50μm distance bin. Relative difference between the ratio of synchronized (black line) to non-synchronized neighbors (blue dotted line) across distance in the network (*righthand axis*; Kolmogorov-Smirnov test, D=0.13, p<0.0001, right).

### Non-hippocampal drive of MEC synchrony

To establish whether functional hippocampal connections to MEC exist *in vitro* during the second postnatal week, we measured field excitatory postsynaptic potentials (fEPSPs) in sMEC evoked by stimulation of dentate gyrus (DG) or CA1 in the immature (P8-P10) hippocampus and compared them with local stimulation in deep layers of MEC (dMEC) (Fig.4a, *left*, 4b,c). Using a 60-electrode array in MEC (Fig. 4a, *left*) we recorded high failure rates for both CA1- and DG-stimulation (76% and 57% of total experiments, respectively) compared with a total absence of failures upon dMEC stimulation (Fig.4c). Therefore, direct functional connections exist between the hippocampus and sMEC but they are highly unreliable and significantly weaker than local projections from dMEC layers.

**Figure 4:**
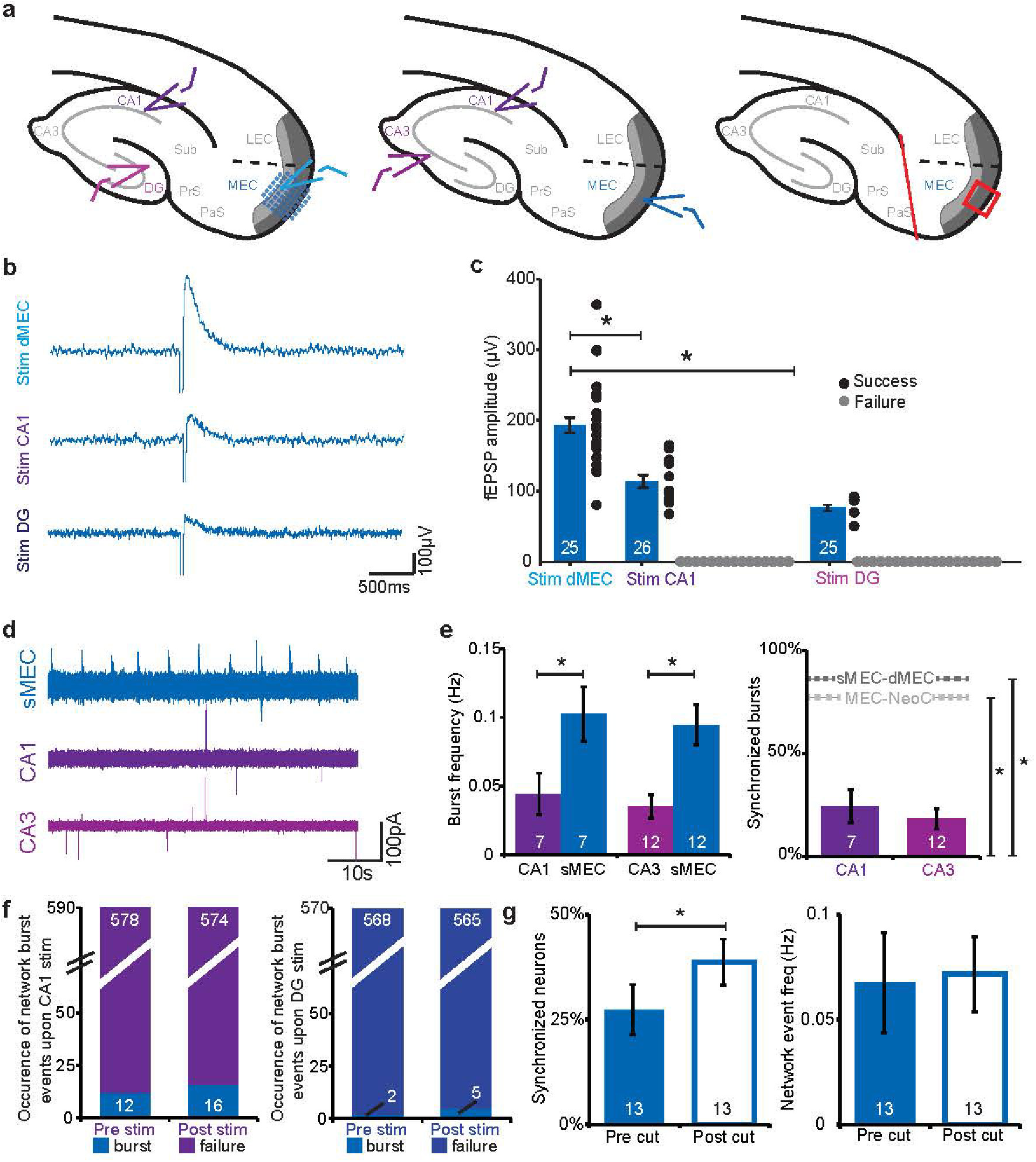
Immature hippocampus does not drive synchronous network activity in MEC. (**a**) *Left*, Overview of MEA recordings (blue grid, MEC) and stimulation sites (Dentate gyrus (DG), CA1 and dMEC), *middle*, field recording sites (CA1, CA3, sMEC) and *right*, recording site of multiphoton network calcium imaging (red box) with hippocampal lesion (red line). (**b**) Example traces of mean evoked fEPSPs of representative experiments (n=20 sweeps) upon dMEC, CA1 and DG stimulation recorded using a multielectrode array in sMEC. (**c**) Evoked fEPSP amplitudes (black) and response failures (gray) for dMEC (n=25 slices, 0% failures), CA1 (n=26 slices, of which 76% failed to evoke sMEC) and DG (n=25 slices, 57% failures). Mean of successful responses was significantly bigger in sMEC than in CA1 (unpaired t-test, t(34)=4.00, p<0.001) and DG (unpaired t-test, t(29)=4.57, p<0.0001). (**d**) Example of typical field recordings (1Hz highpass filtered) show rhythmic burst activity in sMEC but less frequent, single spike activity in CA1 and CA3. (**e**) *Left*, Network event frequency is significantly lower in CA1 and CA3 compared with sMEC (paired t-tests, t(6)=6.95, p<0.001 and t(11)=6.12, p<0.0001 respectively). *Right*, low levels of synchrony exist between CA1-sMEC and CA3-sMEC that are significantly weaker than sMEC-dMEC and sMEC-NeoC (Unpaired t-tests, CA1-sMEC vs. NeoC-sMEC: t(15)=3.87, p<0.01, CA3-sMEC vs. NeoC-sMEC: t(20)=5.72, p<0.0001, CA1-sMEC vs. dMEC-sMEC: t(12)=6.67, p<0.0001 and CA3-sMEC vs. dMEC-sMEC: t(17)=9.24, p<0.0001) gray dashed lines, see Figures 5 and 6 respectively for comparative data. (**f**) Evoked field stimulation in CA1 (*left*) and DG (*right*) failed to reliably evoke network bursts in sMEC and did not alter the rate of spontaneous network bursts (chi-square with Yates correction, CA1: chi-square(1)=0.33, p=0.56, DG: chi-square(1)=0.58, p=0.45). (**g**) Removal of hippocampus significantly increased the number of synchronized neurons (paired t-test, t(7)=3.09, p<0.05) but did not change the network frequency of synchronized events in sMEC (paired t-test, t(7)=1.23, p=0.26). *N experimental numbers indicated on bars*.

To test whether spontaneous network activity in sMEC could be driven by the immature hippocampus we made simultaneous field recordings in CA1-sMEC, CA3-sMEC and CA1-CA3-sMEC (Fig.4a *middle*, Fig.4d). Frequency of spontaneous activity was significantly higher in sMEC compared to CA1 (0.1±0.02Hz vs. 0.04±0.015Hz, Fig.4e) and CA3 (0.09±0.015Hz vs. 0.04±0.008Hz, Fig.4e). Furthermore, the proportion of network bursts that were synchronized with sMEC (Fig.4e, *right*), was significantly lower for CA1 (24±8%) and CA3 (18±5%) than the clear, consistent synchrony observed between sMEC-dMEC (90±5%) and sMEC-Neocortex (NeoC) (79±10%, Fig. 4e, see also Fig.5c).

**Figure 5:**
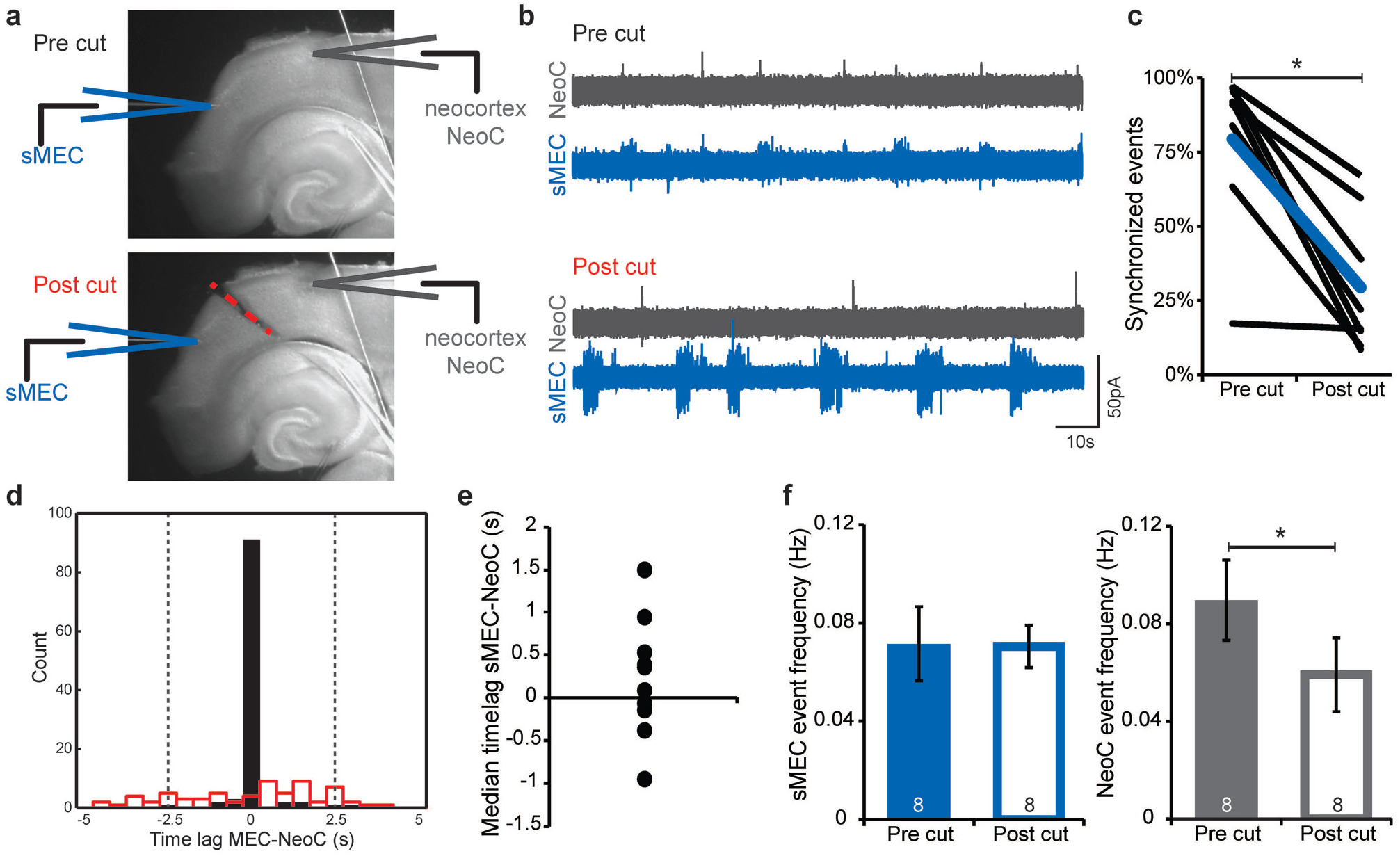
Spontaneous network activity in immature MEC is synchronized with but not driven by neocortex. (**a**) Schematic view of field recordings and lesion site. (**b**) Example recordings of simultaneously pre- (top traces) and post-lesion (lower traces) sMEC (blue) and superficial neocortex (NeoC, gray). (**c**) Significant decrease in the proportion of synchronized events between sMEC and NeoC post-lesion (paired t-test, t(7)=5.10, p<0.01). Blue line indicates mean values, n=8 slices. (**d**) Representative time-lag histogram for onset times of spontaneous network activity between sMEC and NeoC pre- (black bars) and postlesion (red open bars). All events with a timelag ±2.5s (gray dotted lines) are considered synchronized. (**e**) No reliable time-lag (one-sample t-test, t(9)=1.06, p=0.32) indicating no directionality of activity between sMEC and NeoC. (**f**) No change in the frequency of sMEC network activity following lesion (paired t-test, t(7)=0.10, p=0.92) but significant decrease in NeoC network activity (paired t-test, t(7)=3.19, p<0.05). *N experimental numbers indicated on bars*.

To verify that CA1 and CA3 do not initiate MEC events, we quantified the occurrence of bursts of sMEC network activity one second before and after field stimulation in CA1 and DG (Fig.4a *left*, see Methods for ‘network burst’ definition). The majority of sweeps showed no evoked sMEC network burst within one second following stimulation in either CA1 (574/590 trials post-stim, Fig.4f, *left*) or DG (565/570 trials post-stim, Fig.4f, *right*). Furthermore, the low occurrence of network bursts upon stimulation did not differ from spontaneous burst rates observed up to one second prior to stimulation in CA1 (12 events = 2% trials pre-stim, 16 events = 3% trials post-stim) or DG (2 events <1% trials pre-stim, 5 events <1% trials post-stim), indicating that the hippocampus cannot reliably evoke the developmental network bursts in sMEC. Finally, to investigate the modulatory effect of the hippocampus proper upon sMEC network events, we recorded network events using calcium imaging before and after a hippocampal lesion (Fig.4a, *right*). Following removal of hippocampal inputs, the number of synchronized neurons within sMEC increased significantly (27±6% to 39±6%, Fig.4g, *left*). However, the network event frequency remained stable (pre:0.07±0.024, post: 0.07±0.018, Fig. 4g, *right*) demonstrating that the hippocampus modulates but does not drive the synchronized spontaneous sMEC network activity.

### Neocortical synchrony is paced by intrinsic sMEC activity

Immature entorhinal cortex has been proposed as a cortical ‘pacemaker’ whose intrinsic activity drives neighbouring neocortical regions during early postnatal development (Garaschuk, Linn et al. 2000),(Namiki, Norimoto et al. 2013). To firstly confirm that sMEC can self-generate its own network activity, we utilized calcium imaging in isolated MEC mini-slice preparations (Suppl. Fig 3a). Frequency of network events dropped upon isolation (0.12±0.02Hz vs. 0.06±0.01Hz, Suppl. Fig.3c, *right*). However, we observed no significant changes in fractions of active (47±3% vs. 47±5%, Fig.5b, Suppl. Fig 3b, *left*), synchronized neurons (31±4% vs. 29±6%, Suppl. Fig.3b, *right*) or activity levels (0.04±0.009Hz vs. 0.02±0.006Hz, Suppl. Fig.3c, *left*) indicating that intrinsic synchrony persists, similar to the intact slice preparation. To test whether sMEC bursts drive neocortical activity, we then used paired field recordings in both sMEC (blue traces) and NeoC (gray traces) to measure spontaneous network bursts (Fig.5a,b *top panels*). NeoC network bursts were highly synchronized with sMEC bursts (Fig.5b, *top*, Fig.5c 0.79±0.1, ‘pre-cut’,5d). We saw no significant time-lag between sMEC and NeoC to indicate a consistent origin of activity and propagation (230±218ms, Fig.5d: example recording, 5e). To directly test if sMEC paces NeoC activity, we lesioned all interregional connections (Fig.5a, *bottom*). Both sMEC and NeoC displayed rhythmic network bursts following lesioning (Fig.5b, *bottom*), indicating that NeoC activity must be partly generated by a source other than sMEC. However, sMEC-NeoC synchrony was strongly decreased post-lesion (Fig.5c, pre: 79±10%, post: 29±8%, red bins Fig.5d). NeoC network burst frequency significantly dropped following separation from sMEC (0.09±0.02Hz vs. 0.06±0.02Hz, Fig.5f, *right*). In contrast, no significant change in frequency of network bursts occurred in sMEC (0.07±0.02Hz vs. 0.07±0.01Hz, Fig.5f, *left*), suggesting that NeoC bursts synchronize to sMEC activity.

### dMEC-sMEC time-locked network synchrony

Pyramidal neurons in dMEC connect to both principal neurons and interneurons in adult sMEC (van Haeften, Baks-te-Bulte et al. 2003) (Hamam, Kennedy et al. 2000). To assess whether dMEC network events synchronize with sMEC during the second postnatal week of development, we measured their activity simultaneously using low-magnification calcium imaging (Fig.6a). dMEC and sMEC layers were spatially-defined prior to event analysis (Fig.6a). Synchronized neurons occurred in both dMEC and sMEC layers (Fig.6b) with no difference in density between layers or relative to pial distance (Fig.6b). Network events were synchronized across individual neurons in deep and superficial layers (Fig.6c) with a similar rhythmic frequency (Fig.6d, 0.04±0.01Hz vs. 0.03±0.01Hz). For millisecond-level resolution, we investigated the temporal relationship between dMEC and sMEC using paired cell-attached recordings between sMEC stellate cells and dMEC pyramidal neurons and also field recordings (Fig.6e-h, Suppl.Fig.4a-c). Network bursts were highly synchronized (Fig.6g, 76±6%, n=14(cell-attached), Suppl.Fig.4b,c, 89±5%, n=7(field recordings)). There was no significant time-lag between cell pairs or layers, providing no evidence that deep layers were driving spontaneous superficial network activity (Fig.6h, cell-attached median timelag: 6ms IQR: −0.14 to +0.36ms, Suppl. Fig.4c field recordings median timelag: −286ms IQR: −1125 to +161ms). Furthermore, paired whole-cell voltage-clamp recordings from identified stellate cells in sMEC and dMEC pyramidal neurons confirmed strikingly similar spontaneous input patterns during network bursts (Fig.6e,g, lower traces).

**Figure 6:**
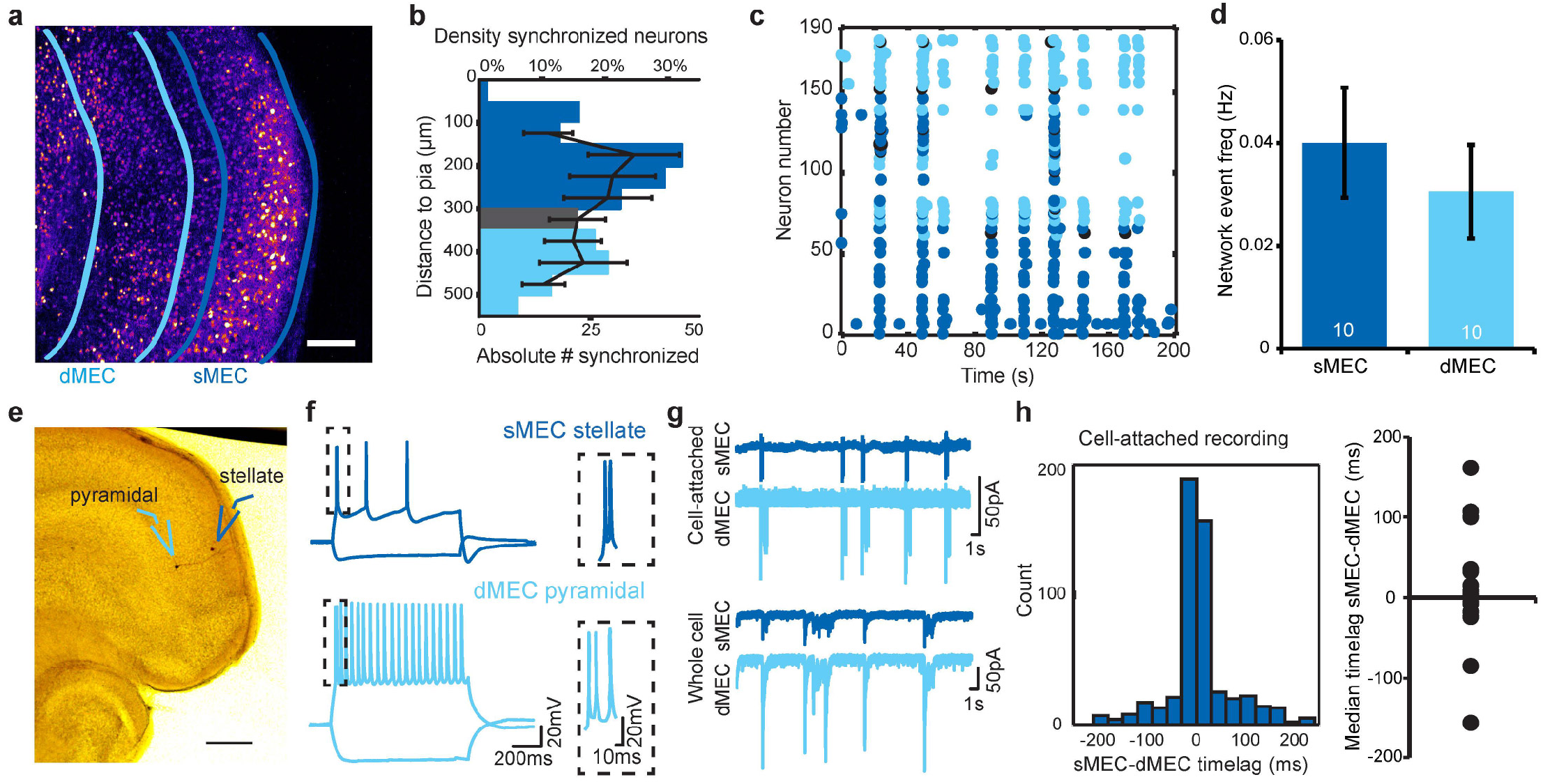
Deep and superficial layer immature MEC networks are synchronized. (**a**) Two-photon image of entire intact MEC loaded with Fura2-AM calcium dye. Example locations for imaging in deep (light blue) and superficial (dark blue) layers indicated. *Scalebar 100μm*. (**b**) Density of synchronized neurons (black lines) is not different between deep and superficial layers (one-way ANOVA, F(7,69)=0.76, p=0.63). Absolute number of synchronized neurons per distance bin indicated as bars. (**c**) Representative raster plot of deep (light blue), superficial (dark blue) and intermediate layer (gray) MEC activity showing correlated synchrony across layers. (**d**) Quantification reveals no significant difference in frequency of synchronized network activity between deep and superficial networks (paired t-test, t(9)=0.98, p=0.35). N numbers indicated (**e**) Representative biocytin-filled spiny stellate and deep layer pyramidal neuron taken from paired recordings in dMEC and sMEC (*left*, scalebar 250μm). (**f**) Cell-attached (n=14 pairs) and (**g**) whole cell recordings (n=6 pairs) showing synchrony of spike rates and synaptic inputs. (**h**) Histogram of the observed time-lag for event onset using high temporal resolution cell-attached recordings show correlated synchrony between deep and superficial layers but no directionality (Wilcoxon Signed Rank test W=64, p=0.52) in immature MEC.

In addition to spontaneous activity, burst activity could be evoked simultaneously and reliably in both superficial layer stellate cells and in layer 5/6 pyramidal neurons, following field stimulation in sMEC (LV/VI neuron: 0.82 ± 0.1, LII stellate cell: 0.9 ± 0.05 Suppl Fig 4d). Interestingly, generation of network burst activity was not possible if preceded by spontaneous network activation seconds before and it was followed by a refractory period of approximately 10 seconds during which the network was not synchronously active (example sMEC stimulation, Suppl. Fig 4e).

Therefore, we found no evidence supporting the idea that dMEC is driving spontaneous synchronous network activity but rather that its activity is synchronized to that of the superficial layers.

### Glutamatergic activity underlies immature MEC synchrony

Synchronized network activity during the first postnatal week is dependent upon glutamatergic or GABAergic mechanisms in NeoC and hippocampus, respectively (Allene, Cattani et al. 2008) (Garaschuk, Hanse et al. 1998, Garaschuk, Linn et al. 2000) (Conhaim, Easton et al. 2011). Given the prominent GABAergic connectivity within adult sMEC (Couey, Witoelar et al. 2013, Pastoll, Solanka et al. 2013) and that MEC is part of the parahippocampal region where early hippocampal network activity is mediated via GABAergic signalling (van Strien, Cappaert et al. 2009), we hypothesized that spontaneous network activity would be mediated by GABAergic signalling. To rule out glutamatergic mechanisms, AMPA and NMDA receptors were selectively blocked (Fig.7a). Blockade of either receptor decreased overall frequency of activity (CNQX: 0.03±0.002Hz vs. 0.02±0.003Hz, DAPV: 0.06±0.005Hz vs. 0.034±0.003Hz, Fig.7b *left*). However, the number of synchronized neurons after application of each blocker did not alter significantly (CNQX: 27±9% vs. 22±9%, DAPV: 46±4% vs. 38±6%, Fig.7b *right*), thus any persistent network activity remained synchronous. The drop in frequency was not due to a subgroup of neurons decreasing their frequency but by all neurons (example slices: Fig.7c, Suppl.Fig.5).

**Figure 7:**
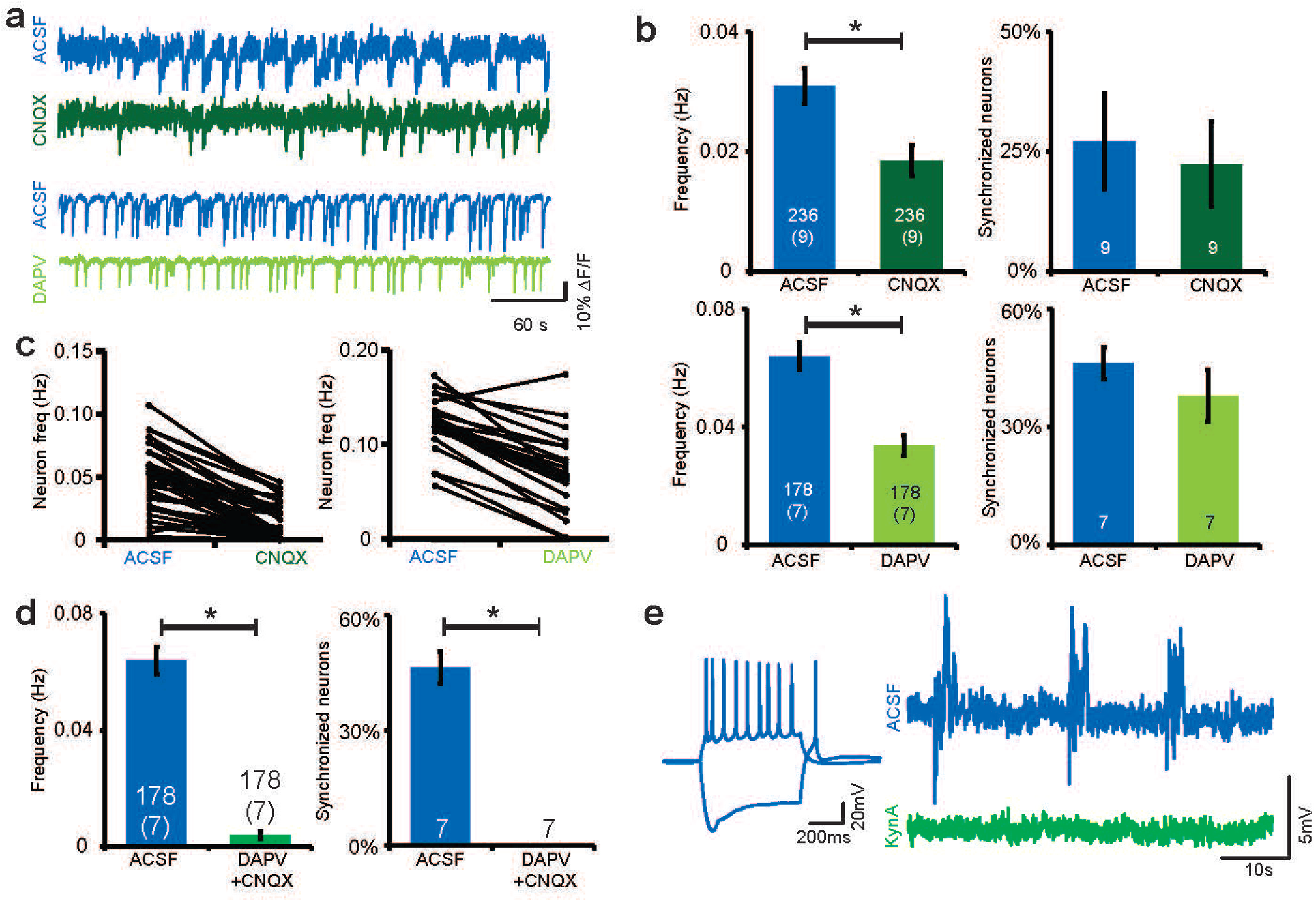
Ionotropic glutamatergic signaling underlies spontaneous synchronized network activity in immature sMEC. (**a**) Example traces of individual neurons before (blue, ACSF) and after inhibition of AMPA (dark green) and NMDA receptors (light green). (**b**) Blockade of AMPA or NMDA receptors decreased overall frequency of activity (paired t-test, CNQX: t(235)=5.93, p<0.0001, DAPV: t(177)=10.98, p<0.0001) but not the proportion of synchronized neurons (paired t-test, CNQX: t(8)=1.07, p=0.31, DAPV: t(6)=1.22, p=0.27). *N*=*9 and 7 slices containing 236 and 178 neurons for CNQX and DAPV experiments respectively*. (**c**) Significant change in frequency occurred across the majority of neurons following application of either ionotropic glutamate receptor blocker. Example experiments shown (CNQX: 2 increasing, 50 decreasing frequency, DAPV: 1 increasing, 40 decreasing frequency). (**d**) Combined blockade of AMPA and NMDA receptors significantly decreased network activity (left) and abolished all synchronous suprathreshold network activity (*right*) in sMEC (paired t-test: t(177)=13.01, p<0.0001 and t(6)=11.22, p<0.0001 respectively). (**e**) Left: Spike profile of developing stellate cell. *Right*: Blockade of ionotropic glutamate receptors by kynurenic acid (green) also abolished subthreshold excitatory synaptic activity from individual stellate cells.

In contrast, simultaneous blockade of both receptor types dramatically reduced the frequency of network activity (0.064±0.005Hz to 0.004±0.001Hz, Fig.7d, *left*) and abolished synchronized suprathreshold activity in all slices (46±4% for ACSF to 0%, Fig.7d, *right*). Whole-cell patch-clamp recordings were made from stellate cells to assess subthreshold membrane potential changes before and after blockade of all ionotropic glutamate receptors. Clear periodic network bursts of synaptic activity resulting in action potentials were observed in ACSF (blue trace, Fig.7e). Kynurenic acid application silenced all rhythmic suprathreshold and subthreshold activity in stellate cells (green trace, Fig.7e, n=4, group data not shown), thus, synchronized network events in sMEC and rhythmic activity of stellate cells were silenced by blockade of ionotropic glutamate receptors at this immature stage, similar to glutamatergic regulation in immature neocortical circuits.

### Non-synaptic mechanisms modulating network activity

Prior to GABAergic and glutamatergic synaptic modulation, neocortical synchrony is modulated by gap junctions perinatally in the neocortex and retina (Sun and Luhmann 2007) (Blankenship and Feller 2010). Therefore, we tested whether non-synaptic mechanisms also affected early sMEC networks. Gap junction blockers selectively reduced frequency of network activity (0.08±0.012Hz vs. 0.03±0.007Hz, Suppl.Fig.6a) but did not change the number of active (57±6% vs. 54±6%) or synchronized neurons (41±7% vs. 34±6%, Suppl.Fig.6a). Thus, gap junctions contribute to immature sMEC synchronous network activity but given the residual synchrony after pharmacological blockade, are not the driving mechanism.

Prolonged electrical bursting activity in pyramidal neurons depends on extracellular calcium concentrations in hippocampus and entorhinal cortex with decreased levels inducing prolonged bursting (Su, Alroy et al. 2001, Sheroziya, von Bohlen und Halbach et al. 2009). Therefore, we tested if changing extracellular calcium levels modulated activity across the sMEC network and further, whether it restored downregulated network activity at the end of the second postnatal week. Indeed, we show that while network composition remained stable, increasing extracellular calcium reduced frequency of activity (0.07±0.006 vs. 0.05±0.004, Suppl. Fig.7a, *right*) while decreasing levels caused an increase of activity levels at P7-12 (0.07±0.006Hz vs. 0.08±0.006Hz, Suppl. Fig.7a, *right*) and enhanced the low activity levels at P13/14 - albeit not restoring it completely to peak activity levels (P7-12) (0.003±0.001Hz vs. 0.010±0.002Hz, Suppl. Fig.7b, *right*). This effect was not mediated by NMDA receptors (Suppl Fig. 7c). Thus, extracellular calcium levels play a role to upregulate suprathreshold network activity but cannot restore synchrony in sMEC at the end of the second postnatal week.

### GABAergic modulation of network activity

From the third postnatal week of development onwards strong feedback inhibition is essential for network function and rhythmic activity in sMEC (Couey, Witoelar et al. 2013, Pastoll, Solanka et al. 2013). Spontaneous network activity in sMEC becomes asynchronous and sparse at the end of the second postnatal week (Fig. 2). We hypothesized that increased GABAergic tone and maturation of the GABAergic system could mediate the abolition of synchronous network activity. Thus, to determine the role of GABA, we tested the blockade of both GABA-A and GABA-B receptors during synchronous activity at the start of the second postnatal week and during asynchronous, sparse activity at P13-15 (Fig.8a). Blockade of GABA-A receptors increased the number of active neurons within the network at both developmental stages (P8: +9±4%, P14/15: +12±6%, Fig.8b, *top left*). Despite an increase in the number of synchronized neurons upon GABA-A blockade (P8/9: +12±7%, P14/15: +18±7%, Fig.8b, *bottom left*), the frequency of network events in these synchronized neurons did not increase (P8/9: −0.01±0.01Hz, P14/15: −0.01±0.02Hz, Fig.8b, *bottom right*). However, GABA-A receptor blockade did cause a significant increase in overall activity at P14/15 but a decrease at P8/9 (P8/9: −0.002±0.001, P14/15: +0.014±0.003, Fig.8b, *top right*) suggesting a developmental increase in an inhibitory effect of GABA-A mediated signaling to modulate network activity.

**Figure 8:**
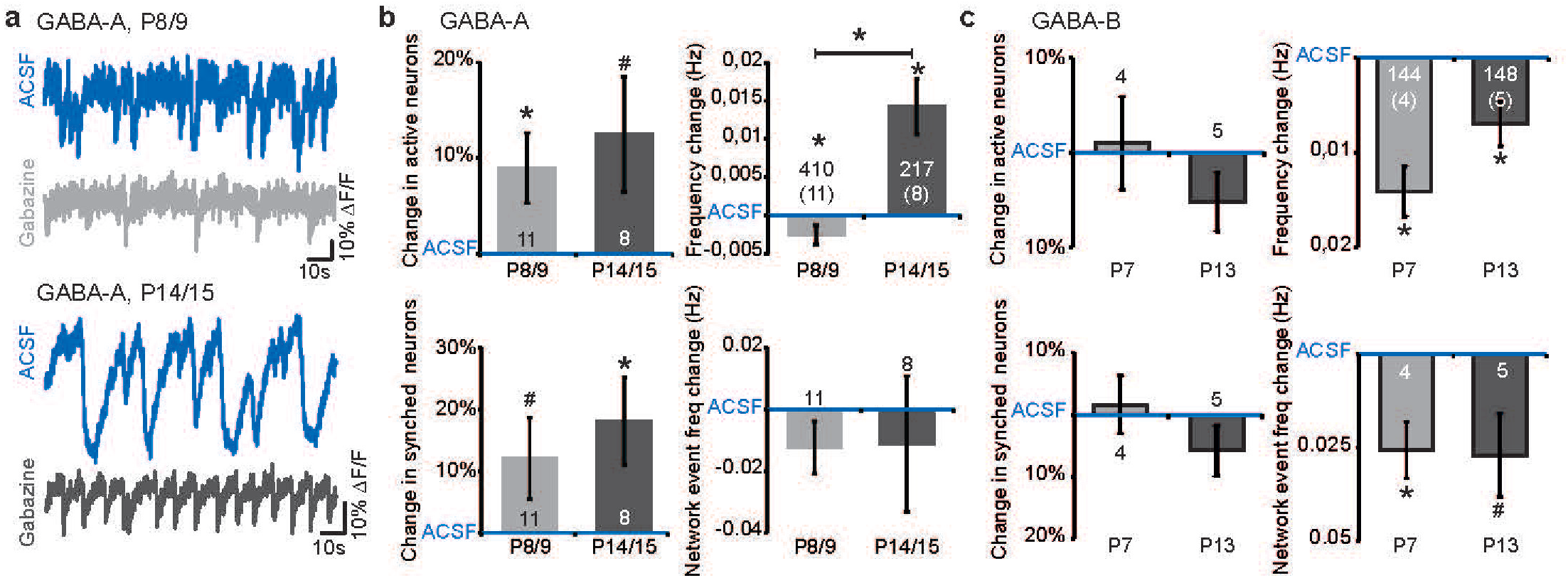
GABA-A and GABA-B receptor signaling modulates but does not drive network activity. (**a**) Example traces before and after blockade of GABA-A receptors at P8/9 group and at P14/15 group. (**b**) Gabazine application increased the amount of active (one-sample t-test, P8/9: t(10)=2.44, p<0.05 and trend at P14/15: t(7)=2.06, p=0.08, *top left*) and synchronized neurons (one-sample t-test, P14/15: t(7)=2.56, p<0.05 and trend at P8/9: t(10)=1.83, p=0.01, *bottom left*). No significant change of network event frequency upon Gabazine application (one-sample t-test, t(10)=-1.45, p=0.18 and t(7)=-0.51, p=0.63 for P8/9 and P14/15, *bottom right*). No age-dependent differences of Gabazine’s effect on active or synchronized neurons and network event frequency (independent t-test, t(17)=-0.519, p=0.61, t(17)=-0.62, p=0.55 and t(17)=-0.05, p=0.97 respectively). Frequency of activity was differentially affected by Gabazine depending on age (independent t-test, age effect: t(625)=-5.36, p<0.0001, one-sample t-test, condition effect P8/9: t(409)=-2.07, p<0.05 and P14/15: (216)=3.94, p<0.001, *top right*). (**c**) GABA-B receptor effects were not developmentally regulated between P7 and P13 (independent t-test: active neurons: t(7)=1.12, p=0.30, frequency: t(290)=-2.04, p=0.48, synchronized neurons: t(7)=1.20, p=0.27, network event freq: t(7)=0.10, p=0.93). No significant effect of GABA-B blockade on proportion of active (one-sample t-test, P7: t(3)=0.21, p=0.85, P13: t(4)=-1.67, p=0.17, *top left*) or synchronized (one-sample t-test, P7: t(3)=0.35, p=0.75, P13: t(4)=-1.42, p=0.23, *bottom left*) neurons. Significant decrease of activity (one-sample t-test, P7: t(143)=-5.35, p<0.0001, p13: t(147)=-2.97, p<0.01, *top right*) and network frequency (one-sample t-test, P7: t(3)=-3.41, p<0.05 and a trend P13: t(4)=-2.4, p=0.07, *bottom right*). *N-numbers indicated in individual bars indicating slices or neurons and slices in brackets*.

In juvenile MEC (third postnatal week), metabotropic GABA-B receptors play a role in terminating persistent network activity of layer III pyramidal neurons (Mann, Kohl et al. 2009). In contrast to GABA-A receptor blockade, GABA-B receptor blockade did not increase the number of active nor synchronous neurons (Fig.8c *left*). However, inhibition of GABA-B receptors decreased the frequency of activity and the rate of synchronous firing at both timepoints measured (P7:-0.03±0.008Hz, P13:-0.03±0.011Hz and P7:-0.1±0.003, P13:-0.007±0.002, Fig.8c *right*). Thus, GABA-A and GABA-B effects are mediated in an opposing manner to each other at this immature stage of development. Importantly, GABA-A and GABA-B receptors do not drive synchrony but significantly modulate network activity levels overall by P13/15 and regulate activity in comparable patterns to those reported from the third postnatal week onwards.

## Discussion

GABAergic inhibition plays a key role in regulating sMEC activity from the third postnatal week of development onwards, when distinct spatially-tuned firing properties of cells first appear *in vivo* (Langston, Ainge et al. 2010, Couey, Witoelar et al. 2013, Pastoll, Solanka et al. 2013, Wills and Cacucci 2014). In contrast, our data show that prior to spatial exploration and eye-opening in the rodent, intrinsic sMEC network activity is highly synchronous both *in vivo* and *in vitro*, and mediated by glutamatergic not GABAergic mechanisms. sMEC synchronous networks occur in a non-random pattern through superficial layers, akin to the functional modules theoretically predicted (McNaughton, Battaglia et al. 2006). Synchronized activity is time-locked across both deep and superficial layers in layer V pyramidal neurons and layer II stellate cells and entrains synchronized activity in neighboring neocortex. These synchronized rhythmic patterns of activity are transient, disappearing as GABAergic inhibition matures by the end of the second postnatal week.

Our data clearly show that synchronous sMEC network activity and rhythmic firing of stellate cells is mediated by glutamatergic excitation during the second postnatal week of development. This network activity contrasts with the predominant dense recurrent GABAergic inhibition one week later and in young adult rodents (Couey, Witoelar et al. 2013, Pastoll, Solanka et al. 2013) that has been shown to be sufficient to generate, for example, grid cell firing patterns in attractor dynamic models (Couey, Witoelar et al. 2013, Pastoll, Solanka et al. 2013). Mature entorhinal cortex has a clear modular organization, with dendritic and axonal clusters of 400-500μm projecting from deeper layers into layer II (Ikeda, Mori et al. 1989, Hevner and Wong-Riley 1992) (Solodkin and Van Hoesen 1996). Models of early MEC development predict topographically-organized modules of both excitatory and inhibitory connectivity that form in an experience-independent manner (McNaughton, Battaglia et al. 2006). However, there is a lack of data determining the relevant biological parameters for computational models of MEC development before recurrent inhibition. Here, our findings reveal a synchronized module of neurons non-randomly distributed throughout sMEC layers up to a maximal distance of 450μm in a developmental period just prior to eye-opening and explorative behavior. In these immature networks, stained synchronized neurons were just over 150μm apart and significantly closer to each other than to their active but asynchronous neighbors. This distance is comparable to the distribution ranges observed for connectivity maps of stellate cells and for the patch-like distribution of cytochrome oxidase staining in rodent sMEC (Burgalossi, Herfst et al. 2011) (Beed, Bendels et al. 2010). Thus, we hypothesize that these highly synchronous immature networks of neurons are an early precursor for later formation of patches of functionally connected neurons.

In the adult, excitatory drive from the hippocampus to sMEC modulates grid cell function (Bonnevie, Dunn et al. 2013) and long-range hippocampal GABAergic projections alter rhythmic network activity in MEC (Melzer, Michael et al. 2012). Connectivity between parahippocampal regions of pre- and parasubiculum to MEC exists by P10 (Canto, Koganezawa et al. 2012). Thus, by P10 MEC receives extrinsic inputs. However, at peak network activity and synchrony ages, we observed only weak and unreliable functional long-range connections from hippocampus proper to immature sMEC. In the adult brain, high levels of synchrony occur between MEC and CA1 hippocampus (Hahn, McFarland et al. 2012) whereas we show that in immature brain, both DG and CA1 were synchronized only approximately 25% of the time with sMEC network bursts compared to 79% synchrony between MEC and neocortical regions at the same age. Further, we demonstrate that in contrast to the driving influence of hippocampus onto adult MEC(Melzer, Michael et al. 2012, Bonnevie, Dunn et al. 2013), stimulating hippocampal fibers directly did not relieably induce bursting activity in immature MEC.

The entorhinal cortex is proposed to act as a pacemaker for cortical Early Network Oscillations (cENOs) during the first postnatal week (Garaschuk, Linn et al. 2000, Namiki, Norimoto et al. 2013). Our results show that MEC is indeed pacing the rate of rhythmic activity in neighboring neocortex during the second postnatal week since lesioning between the two regions caused a decrease, but no disappearance of neocortical activity alone. Further, the high level of synchrony between MEC and neocortex greatly reduced after the lesion. The residual NeoC activity could be intrinsic to the local circuit, potentially entrained by earlier prenatal subplate activity (Hanganu, Okabe et al. 2009).

Due to strong connectivity from deep to superficial MEC (Dhillon and Jones 2000, van Haeften, Baks-te-Bulte et al. 2003, Beed, Bendels et al. 2010) we anticipated that deep layers would drive spontaneous rhythmic activity in superficial networks. However, our data showed strongly time-locked synchrony at both a network level between deep and superficial layers and at individual neuron level between layer V pyramidal and layer II stellate cells. We extend the finding in individual layer III neurons that synchronous activity is intrinsic to immature MEC (Sheroziya, von Bohlen und Halbach et al. 2009) to a network level across multiple MEC layers. Both layer V pyramidals and stellate cells receive synchronous synaptic inputs, similar to the common synaptic inputs on neurons across all layers of MEC from pre- and parasubicular regions at P14-31 (Canto, Koganezawa et al. 2012). We were able to induce network burst activity in both layer V pyramidals and stellate cells following local stimulation in superficial MEC, providing that stimulation did not occur immediately following spontaneous burst generation. For synchronous bursts, the exact cellular origin of this immature activity remains to be determined. In lateral entorhinal cortex during the first postnatal week, superficial neurons drive bursting activity within deeper layers with second-long timelags (Namiki, Norimoto et al. 2013). However, we see no evidence for significant timelags between deep and superficial layers. It remains to be determined whether hub neurons exist in MEC that could coordinate or induce the intrinsic network activity, similar to GABAergic interneurons in developing hippocampus (Bonifazi, Goldin et al. 2009).In support, stellate cells in sMEC, on the basis of maturational markers and time-specific neuronal silencing, are proposed as the source that drives maturation of the entorhinal cortex-hippocampal network (Donato, Jacobsen et al. 2017).

GABAergic inhibition mediates rhythmic synchronized network activity within immature rodent cerebellum (P4-6, (Watt, Cuntz et al. 2009)) and hippocampal CA3 and CA1 regions, categorized in hippocampus as giant depolarizing potentials (GDPs)(P3-12, (Ben-Ari, Cherubini et al. 1989, Garaschuk, Hanse et al. 1998, Khazipov, Khalilov et al. 2004)). In contrast to the hippocampus, glutamatergic network mechanisms and synchrony in immature sMEC are similar to those underlying early synchrony throughout neocortex during the first postnatal week (cENOs) (Garaschuk, Linn et al. 2000) (Namiki, Norimoto et al. 2013) (Conhaim, Easton et al. 2011). The loss of spontaneous network activity upon inhibition of AMPA and NMDA receptors could be explained by a loss of excitatory input to GABAergic interneurons that potentially drive activity within the network. However, blocking GABA-A receptors directly did not decrease the fraction of active nor synchronized neurons within either age group tested but rather increased the numbers of active and synchronized neurons. Thus, despite the strong reciprocal connectivity of parahippocampal regions, including MEC itself, to the hippocampus, distinct mechanisms underlie immature synchrony in these circuits.

We find that suprathreshold network synchrony of MEC is tightly developmentally regulated and transient. Peak measures of synchronous activity and maximal levels of synchronized neurons at P10/11 correspond to those observed in somatosensory and visual cortices prior to eye-opening in rodents *in vivo* (Golshani, Goncalves et al. 2009) (Siegel, Heimel et al. 2012) (Rochefort, Garaschuk et al. 2009). What curtails suprathreshold network activity within MEC? By the end of the second postnatal week, we observed sparse and largely asynchronous sMEC network activity that is modulated by GABA-A receptor blockade - opposite to that observed one week earlier. Although inhibition of GABA-A receptors and altered extracellular calcium waves can upregulate network activity, in agreement with recordings from individual neurons in MEC and neocortex (Sheroziya, von Bohlen und Halbach et al. 2009) (Conhaim, Easton et al. 2011), neither was able to restore synchronized activity across the network. This inhibition of network activity by P14/15 concurs with an upregulated inhibitory network reported in sMEC from the third postnatal week onwards in rodents and reflects functional circuitry in the adult (Couey, Witoelar et al. 2013).

What is the function of early synchronous activity in MEC? Synchronous activity is correlated with synapse maturation and sensory map formation (Blankenship and Feller 2010) and disruption causes aberrant maturation of circuits. (Grubb, Rossi et al. 2003, McLaughlin, Torborg et al. 2003, Mrsic-Flogel, Hofer et al. 2005). Thus, we postulate that rhythmic waves of MEC activity play a key role in experience-independent maturation of local circuitry, entraining the network for the abrupt appearance of spatially-modulated cell firing during the third postnatal week (Wills, Barry et al. 2012). Furthermore, we speculate that the early synchronously active modules we describe may be predecessors of the cross-layer, anatomically-overlapping functional clusters of grid cells in adult (Hafting, Fyhn et al. 2005, Stensola, Stensola et al. 2012).

In summary, our data demonstrate that topographic modules do exist in immature MEC, as predicted by computational modelling. Contrary to our hypotheses, this synchronous activity is intrinsic to MEC and mediated primarily via glutamatergic, not GABAergic activity. We suggest these data have relevance for our understanding of early entorhinal cortex development and are important biological parameters for computational models of spatial processing in the rodent brain.

## Materials and methods

### Animal usage

All *in vivo* experiments were performed in compliance with the German laws and the guidelines of the European Community for the use of animals in research and were approved by the local ethical committee. Pregnant Wistar rats were obtained at 14-17 days of gestation from the animal facility of the University Medical Center Hamburg-Eppendorf, housed individually in breeding cages with a 12h light/12h dark cycle and fed ad libitum. All *in vitro* procedures involving animals were conducted in accordance with Dutch regulations and were approved by the animal ethics committee (DEC) of the VU University Amsterdam. C57BL/6 mouse pups of both sexes aged from postnatal day 7 (P7) to P15 were used for slice experiments.

### Surgical procedure for *in vivo* recordings

Extracellular recordings were performed in the MEC (55mm posterior to bregma and 7.5mm from the midline) using experimental protocols as described previously (Hanganu et al., 2006; Brockmann et al., 2011). Under light urethane-anesthesia (0.125-1g/kg; Sigma-Aldrich), the head of the pup was fixed in the stereotaxic apparatus (Stoelting, Wood Dale, IL) using two metal tubes cemented on to the nasal and occipital bones, respectively. The bone over the MEC was carefully removed by drilling holes of less than 0.5mm in diameter. Removal of the underlying dura mater by drilling was avoided, since leakage of cerebrospinal fluid or blood can dampen the cortical activity and single neuronal firing (I. Hanganu-Opatz, personal observations). The body of the animals was surrounded by cotton wool and kept at a constant temperature of 37°C by placing it on a heating blanket. During recordings, urethane anesthesia (0.1-0.2 times the initial dose) was given when the pups showed any sign of distress. After a 20-60min recovery period, multielectrode arrays (Silicon Michigan probes, NeuroNexus Technologies) were inserted at 10° from the vertical plane into MEC at a depth of 4-5mm. The electrodes were labeled with DiI (1,1’-dioctadecyl-3,3,3’,3’-tetramethyl indocarbocyanine, Invitrogen) to enable post-mortem reconstruction of electrode tracks in the MEC in histological sections (Fig.1a). One or two silver wires were inserted into the cerebellum and served as ground and reference electrodes. Miniature earphones placed under the pup’s body were sensitive enough to detect the smallest visible movements of the limbs as well as the breathing of pups during recordings. The sleep-like conditions mimicked by the urethane anesthesia (Clement, Richard et al. 2008) had no major effect on the properties and dynamics of early network oscillations (Yang, Hanganu-Opatz et al. 2009).

### Protocols for *in vivo* recordings

Simultaneous recordings of LFP and MUA were performed from the MEC using one-shank 16-channel Michigan electrodes (0.5-3ΜΩ). The recording sites were separated by 50μm. The recording sites covered the MEC and lateral entorhinal cortex. Both LFP and MUA were recorded for at least 900s at a sampling rate of 32kHz using a multi-channel extracellular amplifier (Digital Lynx 4S with no gain, Neuralynx, Bozeman, MO) and the corresponding acquisition software (Cheetah). During recording the signal was band-pass filtered between 0.1Hz and 5kHz (Digital Lynx).

### Confirmation of recording locations

DiI (1,1’-dioctadecyl-3,3,3’,3’-tetramethyl indocarbocyanine, Invitrogen) was used for marking electrode recording location *in vivo*. After recording, the pups were deeply anesthetized with 10% ketamine (ani-Medica, Senden-Bösensell, Germany) and 2% xylazine (WDT, Garbsen, Germany) in NaCl (10ml/g body weight) and perfused transcardially with 4% paraformaldehyde dissolved in 0.1M phosphate buffer, pH 7.4. The brains were removed and postfixed in the same solution for at least 24h. Blocks of tissue containing the MEC were sectioned in the coronal plane at 100μm and air dried. Fluorescent Nissl staining was performed using the NeuroTrace^®^ 500/525 green fluorescent Nissl stain (Invitrogen). Briefly, rehydrated slices were incubated for 20min with 1:100 diluted NeuroTrace. Sections were washed, coverslipped with Fluoromount and examined using the green filter (AF 488).

### Analysis of *in vivo* electrophysiology data

Data were imported and analyzed off-line using custom-written tools in Matlab^®^ (Mathworks) software. To detect oscillatory events, the raw data were filtered between 4 and 100 Hz using a Butterworth 3-order filter. Only discontinuous events lasting >100ms, containing at least 3 cycles, and being not correlated with movements (twitches) were considered for analysis. The discontinuous theta bursts in the MEC were analyzed in their occurrence (defined in Hz), duration, max amplitude (defined as the voltage difference between the maximal positive and negative peaks), and dominant frequency. Time-frequency plots were calculated by transforming the LFP using Morlet continuous wavelet. Minimal and maximal intensities in power were normalized to values between 0 and 1 and were displayed in dark blue and red, respectively.

### Preparation of horizontal brain slices

Horizontal entorhinal cortex slices were prepared as described previously (Dawitz, Kroon et al. 2011). Briefly, animals were rapidly decapitated and their brains dissected out under ice cold cutting solution containing (in mM): 110 Choline chloride, 26 NaHCO_3_, 10 D-glucose,11.6 sodium ascorbate, 7 MgCl_2_, 3.1 sodium pyruvate, 2.5 KCl, 1.25 NaH_2_PO_4_ (Merck), and 0.5 CaCl_2_. 300μm thick slices were obtained using a HR2 slicer (Slicer HR-2; Sigmann Elektronik, Hueffenhardt, Germany; vibration frequency: 36Hz, vibration amplitude: 0.7mm, propagation speed: 0.05mm/s). After a minimum recovery period of one hour slices were transferred into a holding chamber containing artificial cerebral spinal fluid (ACSF) with slightly elevated magnesium levels at room temperature composed of (mM): 125 NaCl, 26 NaHCO_3_, 10 D-glucose, 3 KCl, 2.5 MgCl_2_, 1.6 CaCl_2_, and 1.25 NaH_2_PO_4_ (Merck) and continuously bubbled with carbogen gas (95% O2, 5% CO2).

### Reagents

If not indicated otherwise, all reagents were purchased from Sigma-Aldrich. For pharmacology the following concentrations of blockers were added to the standard ACSF: DL-2-Amino-5-phosphonopentanoic acid (DL-APV, Abcam): 100μM, 6-Cyano-7- nitroquinoxaline-2,3-dione (CNQX, Abcam): 2μM, Kynurenic Acid (KynA): 2mM, Gabazine (Tocris): 10μM (blockade of phasic and tonic), CGP 55845 (CGP, Tocris): 4μM, Carbenoxolone (CNX, Tocris): 100μM, and Glycyrrhizic acid (GZA, Abcam): 100μM. Low and high calcium concentrations were achieved by using 0.6mM or 2.6mM CaCl_2_ for the standard ACSF.

### Fura2-AM bulk loading and two-photon data acquisition

After recovery slices were transferred into an interface-chamber filled with 1ml elevated magnesium ACSF heated to approximately 34°C. 50μg fura2-AM (Invitrogen) diluted in 9μl DMSO and 1μl Pluronic^®^ F-127 (20% solution in DMSO; Invitrogen) was pipetted directly on top of the entorhinal cortex and incubated for 20 minutes (for slices of P7/8 animals) to 40 minutes (for slices of P14/15). Slices from animals older than P12 were pre-incubated for three minutes in 3ml ACSF with 8μl 0.5% cremophor (Fluka) diluted in DMSO to facilitate Fura2-AM uptake. After incubation slices were briefly transferred back into the holding chamber and any residual surface dye was washed off.

To improve stability of recordings a poly(ethyleneimine)-solution (1ml poly(ethyleneimine) in 250ml boric buffer containing 40mM boric acid and 10mM sodium tetraborate decahydrate) was used to attach slices onto the recording chambers. After coating of the recording chambers for one hour slices were mounted in elevated magnesium ACSF. The attached slices were placed into a humidified interface container perfused with carbogen and left for at least one hour to achieve a stable attachment and allow for esterase activity to take place within the neurons to trap and activate Fura2-AM. Full methodology details of slice preparation and imaging are published previously (Dawitz, Kroon et al. 2011).

Functional multiphoton calcium dye network imaging data was acquired on a Trimscope (LaVision Biotec) connected to an Olympus microscope using a Ti-sapphire (Coherent) laser tuned to 820nm wavelength. All recordings of sMEC were acquired using a 20x lens (NA 0.95) and a 350x350μm field of view. Parallel recordings of dMEC and sMEC were obtained with a 10x objective (NA 0.3) and a 700x700μm field of view. During data acquisition slices were continuously perfused with oxygenated standard ACSF (recipe as above but 1.5mM MgCl_2_) heated to approximately 27°C. Superficial layers of the medial entorhinal cortex were visually identified using light microscopy. Using a Hamamatsu C9100 EM-CCD camera as a detector, two time-lapse movies (2000 frames each) in sMEC or sMEC and dMEC were acquired with a sampling frequency of approximately 10Hz. For neuron detection, at the end of each recording condition a z-stack ±20μm around the central plane with 1μm slice thickness was acquired.

For simultaneous network imaging and cell-attached stellate cell recording, data was acquired on a Leica RS2 two-photon laser-scanning microscope tuned to 820nm wavelength with a 20X lens and 400x 400μm field of view. Time-lapse data was acquired on a PMT at a rate of 565msec/frame (425 frames each) in sMEC with simultaneous cell-attached recording. Spike rates were acquired using Clampex 10.2 (MDS Analytical Technologies) at an acquisition rate of 10kHz. Cell identity was electrophysiologically- determined with a step profile following breakthrough (see **Patch-clamp recordings** below). Identified stellate cells in layer II were filled with Alexa 594 (80μM, Invitrogen) for additional anatomical identification.

Drugs were washed in for minimally 10 minutes prior to recording except for CNX and GZA that were incubated for at least 30 minutes. Calcium concentrations were changed at least 20 minutes prior to recording onset. Hippocampal and NeoC lesions as well as mini-slices were prepared under a low magnification microscope (4x) using a surgical knife. After lesioning two time-lapse movies and a z-stack were recorded as described above.

### Analysis of imaging data

Custom-built Matlab^®^ (Mathworks) software was used to analyze calcium data (Hjorth, Dawitz et al. 2016). Neurons were detected using the z-stack recorded at the end of each condition. After localizing putative neuronal centers using local intensity peaks, deformed spheres were placed around these centers within the 3D stack to fit the putative neurons. A neuron was added to the contour mask if the spheres had the volume corresponding to a radius of 2-20μm and the intersection of the sphere and the imaging plane had a minimum area of 25μm^2^. The pia was indicated on the final mask to determine the distance to pia of individual neurons in the slice. For analysis of spatial distributions in superficial networks, only neurons located between 50 and 300μm from the pia were analyzed due to sparse cell density in layers I and IV (Fig.3b, gray bars). For low resolution recordings, neurons with 100 to 500μm distance to pia were analyzed.

For each detected neuron the corresponding fluorescent raw trace was extracted from the two consecutive time-lapse recordings and the relative fluorescence trace was calculated (ΔF/F). The baseline was estimated using the running median of the relative trace. Frames with a drop in intensity of at least 10% in the relative trace were considered a putative event. Additionally the event had to be at least 1 SD below the baseline and the trace had to remain significantly below this baseline for a minimum of 5 frames (tested with a one-sided t-test). To confirm onset times and to increase sensitivity of onset detection we repeated this procedure three times excluding the detected putative events from the running median and using a two sided t-test comparing the putative event frames against the filtered baseline for significance in the consecutive iterations. The result was manually inspected and where necessary, corrected.

Groups of synchronized neurons were detected as follows: Onset times of detected events from all neurons were summed together on a frame-by-frame basis and the resulting activity vector was smoothed using a Gaussian. All local peaks exceeding the threshold were defined as network events. The threshold was 5 times the SD of 500 activity vectors derived from the same traces but with randomly shuffled inter-event intervals. A neuron was assigned to the synchronized group if the network events smoothed with a Gaussian overlapped for at least 40% with the total number of event onsets in an individual neuron smoothed with the same Gaussian.

Thus, from this analysis three categories of neurons were derived: silent neurons that do not show any activity, active neurons that show activity and synchronized neurons that are active and whose activity is synchronized to other neurons in the network. The frequency of each group and the network event frequency were calculated. Finally, another Matlab^®^ (Mathworks) script was used to align masks of different experimental conditions to extract repeated measurements of parameters for testing based on individual neurons. All frequency values in the text and figures are derived from these within neuron comparisons and are thus analyzed on single neuron bases, except for the mini-slice experiments (Suppl. Fig 3 where re-alignment of individual neurons was impossible. Here, the average frequencies of all neurons per slice were used. To assess the spatial spread of synchronized neurons within the field of view the average distribution of synchronous neurons around a synchronous neuron was calculated, and compared to the average distribution of neighboring asynchronous neurons. As an additional test, the synchronous neuron distribution was also compared to a shuffled control.

Synchronized activity is referred to as network events to clarify activity measures taken from calcium imaging data. Similar rhythmic activity measured *iIn vivo* and *in vitro* with field and patch-clamp recordings from individual cells are referred to as network bursts throughout the manuscript.

### Field recordings and analysis

dMEC, sMEC, NeoC (perirhinal cortex) and the different hippocampal areas were visually identified with an Olympus microscope (4x lens) using oblique contrast. Field electrodes (chloride-coated silver electrodes inserted into 2-3MΩ borosilicate glass pipettes filled with ACSF) were lowered into the regions of interest using micromanipulators (Luigs-Neuman, Ratingen) and placed at regions with good signal:noise readings. Field potentials were acquired with Multiclamp 400B amplifiers (Molecular Devices) using Clampex 10.2 (MDS Analytical Technologies) at an acquisition rate of 10kHz.

Analysis was carried out using custom-made Matlab^®^ (Mathworks) software. First, signals were high-pass filtered at 1Hz to eliminate any slow frequency drift of the signal. Then, a threshold of 5.5 times the standard deviation (SD) was applied to detect field potentials. For the dMEC-sMEC recordings customized thresholds (between 4.5 and 12 times SD) were used to achieve the optimal detection rates. Spikes were grouped as one event if they were preceded by another spike within 3 seconds. The first spike of an event was defined as the event onset and used to calculate event frequency. Given the long duration of network bursts, a synchronous event between different regions and sMEC was defined as the occurrence of an event in the region of interest within a time-window of 2.5s preceding or following an sMEC event. The proportion of synchrony between different regions relative to sMEC was then calculated.

### Multi-electrode array recordings and analysis

Slices were mounted (see bulk-loading calcium dye methods above) on planar 100/10-ITO-PR multi-electrode arrays (MEAs) consisting of 60 recording electrodes spaced 100μm (Multi Channel Systems, GmbH) with the MEC region positioned on top of the electrode grid (Fig.4).

Spontaneous activity and local field potentials were measured while the recording chamber was constantly bathed with standard ACSF at a flow rate of 4 ml/min and kept at 30±0.5°C. Data acquisition was performed at 5 kHz with MCRack software (Multi Channel Systems). Extracellular stimulation was applied with a silver electrode inserted into a glass borosilicate pipette filled with standard ACSF.

Data was analyzed with custom-made Matlab^®^ (Mathworks) software according to the anatomical layers of MEC. Evoked excitatory post-synaptic potentials were averaged over 20 repetitions and their amplitude was analyzed. For analysis of the occurrence of network burst events prior to and after stimulation, the optimal electrode recordings with sufficient signal to noise ratio to unambiguously detect network burst events were selected in a slice and network burst events up to 1s prior to and 1s after stimulation were quantified.

### Patch-clamp recordings

For patch-clamp recordings the same recording apparatus was used as for field recordings (see above) but the microscope was equipped with a 60x lens. Borosilicate pipettes with a resistance of ∼10MΩ were filled with intracellular solution containing (in mM): 148 K-gluconate, 1 KCl, 10 Hepes, 4 Mg-ATP, 4 K_2_-phosphpcreatine, 0.4 GTP and 0.2% biocytin, adjusted with KOH to pH 7.3. For cell-attached recordings (2-10mins) active cells of interest were selected in dMEC and sMEC and recorded simultaneously in GΩ-resistance seal configurations. After up to 10min continuous voltage-clamp recordings, whole-cell access was made if possible and step protocols were acquired in whole-cell Current-clamp mode to electrophysiologically-identify neuron type and to fill neurons with biocytin for post-hoc anatomical confirmation. To measure synaptic inputs of identified neurons in some cases up to 10min Voltage-Clamp, whole cell recordings were acquired. After recordings, slices were fixed in 4% Paraformaldehyde for at least one week before staining.

Cell-attached data was analyzed using Matlab^®^ (Mathworks) scripts adapted from the field recording analysis. Due to stable baselines, no filtering was applied. Customized thresholds for action-potential (AP) detection ranged from 2-40 times SD. APs were grouped to be part of one network burst if they were preceded by another spike within a timewindow of 1.5s. As a measure of synchrony, the proportion of events in sMEC that are followed or preceded within 1s from time of event onset in the dMEC was calculated.

For anatomical identification slices were stained for biocytin using a modified avidin–biotin– peroxidase method. Note that staining of the entire dendritic tree was limited due to the high input resistance due to the 10ΜΩ pipettes.

### Statistics

For in vivo recordings, data in the text are presented as mean ± SEM. Statistical tests used are indicated in the figure legends for individual experiments. In general, to compare two conditions within the same experiment a paired t-test was used. To compare means from different experiments (i.e. the synchrony or the different ages where GABA-Rs were tested) independent t-tests were performed. Changes upon GABA-R blockade and timelags were assessed based on one-sample t-tests against a hypothetical mean of 0. The difference in mean distance between synchronized to synchronized and synchronized to other neighbors was tested with a Wilcoxon Signed Rank test. The distributions of synchronized to synchronized neighbors and synchronized to non-synchronized neighbors were compared using a Kolgomorov-Smirnoff test.

Multiple means were compared using one-way ANOVAs. Post-hoc analysis was carried out using Bonferroni corrected t-tests. Differences between distributions were assessed using the Kolmogorov-Smirnov test (Fig.3, spatial distributions data). Categorical data (failures and bursts upon stimulation) were analyzed using chi square statistics. Reported is the abbreviation of test statistic with the degrees of freedom in brackets equaling the value of the test statistic. Differences were defined as significant if p-values were smaller than 0.05. A * indicates significance (p<0.05), a # indicates a trend (p<0.01).

## Acknowledgements

This work was supported by the Nederlandse Organisatie voor Wetenschappelijke-Onderzoek (NWO-ZonMW, grant number 917.10.372 to RM), by the European Commission Seventh Framework Programme grant agreement (grant number FP7-People-ITN-2008- 238055 ‘BrainTrain’ to RM) and by Emmy Noether Program of the German Research Foundation (grant numbers Ha4466/3-1, SFB 936 and SPP 1665 to IHO).

